# Computational investigation of biological and technical variability in high throughput phenotyping and cell line identification

**DOI:** 10.1101/175703

**Authors:** Samuel H. Friedman, Paul Macklin

**Author notes:** Twitter: @MathCancer (Paul Macklin).

## Abstract

High-throughput cell profiling experiments are characterizing cell phenotype under a broad variety of microenvironmental and therapeutic conditions. However, biological and technical variability are contributing to wide ranges of reported parameter values, even for standard cell lines grown in identical conditions. In this paper, we develop a mathematical model of cell proliferation assays that account for biological and technical variability and limitations of the experimental platforms, including (1) cell confluency effects, (2) biological variability and technical errors in pipetting, (3) biological variability in proliferation characteristics, (4) technical variability and uncertainty in measurement timing, (5) cell counting errors, and (6) the impact of limited temporal sampling. We use this model to create synthetic datasets with growth rates and measurement times typical of cancer cell cultures, and investigate the impact of the initial cell seeding density and the common practice of fitting exponential growth curves to three cell count measurements. We find that the combined sources of variability mask the sub-exponential growth characteristics of the synthetic datasets, and that researchers profiling the same cell lines under different seeding characteristics can find significant (*p* < 0.05) differences in the measured growth rates. Even seeding the cells at 1% of the confluent limit can cause significant (*p* < 0.05) differences in the measured growth rate from the ground truth. We explored the effect of reducing errors in each part of the virtual experimental system, and found the best improvements from reducing timing errors, reducing cell counting errors, or reducing the interval between measurements (to reduce the inaccuracy of the exponential growth assumption when fitting curves). Reducing biological variability and pipetting errors had the least impact, because any improvements are still masked by cell counting errors. We close with a discussion of recommended practices for high-throughput cell phenotyping and cell line identification systems.

## 1. Introduction and Background

As high-throughput experimental systems advance, novel experiments are using them to measure key cell behaviors (e.g., migration, proliferation, death, and metabolic activity) across a broad range of microenvironmental conditions (e.g., [1-5]). To handle these multicellular phenotypic data and interface them with statistical, mathematical, and computational models, projects such as MultiCellDS [6], PharmML [7], and the Cell Behavior Ontology [8] are developing the consistent data models necessary to analyze, compare, and share experimental data. In particular, the MultiCellDS project is developing a public library of *digital cell lines* that currently aggregate prior cell phenotypic measurements across a variety of microenvironmental conditions. Both original high-throughput experiments and data curation activities require quality control. In experiments, we must determine whether observed differences in cell behavior can be attributed to experimental variability, rather than actual biological differences. When developing community-curated repositories, we must assess whether a submitted measurement is significantly better or worse than an existing measurement. As experimental and informatics platforms merge, we envision systems that seek to match new cell measurements against repositories of known values (e.g., to identify an unknown breast cell sample as behaving more like relatively benign MCF-10A cells or aggressive MDA-MB-231 cells). This, too, will require quantitating the difference between the sample’s measurements and those in the repository.

It is essential for quantification that we assess the impact of experimental design choices, biological and technical variability, instrumentation limitations, and other experimental design choices on the quality of measured parameters. Ideally, we can evaluate a proposed experimental protocol (using a specific set of instruments and analysis software) by comparing its measurements for known cell lines against accepted “gold standard” values as ground truth. However, considerable variability in phenotypic measurements has been observed even for widely used cell lines (e.g., see the discordant cell population doubling times for MCF-7 breast cancer lines grown in “standard” normoxic conditions in [9-17].) Thus, it is difficult to assess the impact of experimental design choices and evaluate the quality of new measurements without well-established ground truths for experimental measurements.

Mathematical modeling can play a role in addressing these experimental challenges by creating idealized systems with fully known ground truths. This approach is frequently used to evaluate new data analysis software: we can evaluate a new algorithm by (1) using a mathematical model to create synthetic datasets with prescribed noise, (2) running the synthetic data through the new tool to estimate cell line properties, and (3) comparing the estimated cell parameters against the original values (the ground truth) (e.g., as in [18]). Recently, mathematicians have used computational models to test the limits of experimental assays, as well as to help us better interpret the biological meaning and significance of new measurement types [19]. Poleszczuk et al. used mathematical models to simulate the spatial distribution of sub-clones in heterogeneous tumors, and analyzed this model system to better understand the impact of where and how we perform tumor biopsies [20]. Treloar et al. fitted mathematical models to several motility assays to demonstrate that assay geometry can have a significant impact on estimated cell proliferation and motility parameters [21]. More recently, Harrison and Baker used mathematical modeling to investigate the impact of temporal sampling limits when measuring cell motility [22].

In this paper, we used a similar approach to investigate the impact of experimental variability and design in estimating cell proliferation rates, as well the role of variability in recognizing when exponential growth models are unsuitable for fitting. Starting with a logistic model of population growth, we developed a technique to create synthetic experimental replicates which account for (1) cell confluency effects in cell culture systems, (2) biological variability in the underlying proliferation parameters, (3) technical variability in micropipetting along with biological variability in cell plating success, (4) technical errors in measurement times along with the impact of sampling limits, (5) technical errors in cell counting accuracy, and (6) technical errors in not accounting for measurement time uncertainties. We used the widespread practice of fitting exponential growth curves to the synthetic data, and noted where experimental variability would be expected to mask the differences between exponential and non-exponential growth.

For a variety of seeding conditions (the number of pipetted cells, as a fraction of the maximum confluent cell population) and experimental variability, we found that the synthetic data would be very difficult to distinguish from exponential growth kinetics, even when analytically we would expect confluency effects to slow growth below exponential. In these cases, experimentalists have justification to fit exponential growth curves to the data and perform standard statistical tests on the differences in estimated growth rates. This can lead investigators to wrongly conclude that the cells had significantly different growth characteristics (*p* < 0.05), even though they actually performed experiments on identical cells. In the scenario where an unknown cell sample is tested against a library of known growth values, an investigator could plausibly but incorrectly conclude that the sample does not match any in the library, even if the library contained the same cell line. We close with an examination of which sources of variability contribute most to misinterpreting data as exponential, and we discuss lessons on improving the robustness of existing experimental protocols.

## 2. Results

We created a simplified digital cell line with a ground truth proliferation rate *α* = 1/30 hr^-1^ ~ 0.033 hr^-1^, giving a mean cell division time of 30 hours and a population doubling time of ~20.8 hours; this value is comparable to many human cancer cell lines [6]. To emulate a physical experimental system (e.g., a well on a multi-well plate or a culture dish), we assigned a fixed population carrying capacity *K* for all numerical experiments. Because we scale the cell counts by *K* and expressed as fractions of the maximum cell capacity, the actual value of *K* does not impact the following results.

Using the ground truth (*α, K*), we then generated synthetic experimental data (See **Methods** below) to investigate the impact of the initial the initial cell seeding density *N*_0_/*K*. For each level of N0/K, we generated 5 biological replicates, and for each biological replicates we generated 20 technical replicates. Each replicate was sampled at 3 times (1, 2, and 3 days after plating) to give synthetic time-course measurements. We used a relatively modest value (5%) for all biological and technical variability and error, and we assumed that all timing errors (the discrepancy between the intended measurement time and its actual measurement time) was normally distributed with a 1 hour standard deviation. See **Methods**.

In Fig. 1, we plotted the synthetic data for each level of the experiment. Note that we used a logarithmic scale, where exponential growth data will appear as straight lines.

**Fig. 1:**
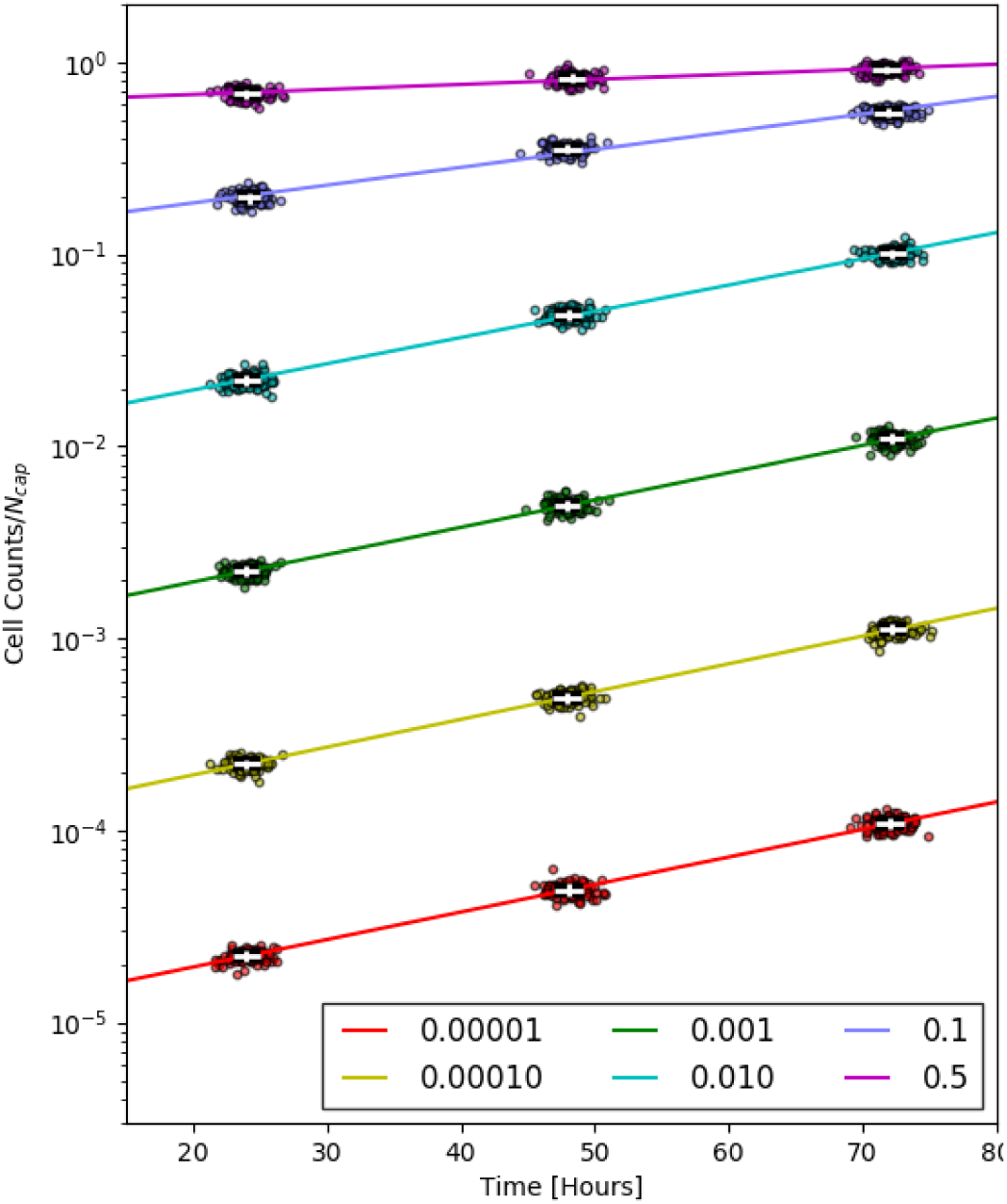
For initial seeding conditions ranging from 10^-5^*K* to 0.5*K* (where *K* is the maximum or confluent cell population), we generated synthetic datasets that simulated biological and technical variability in seeding, growth, and measurements.

As first observation, we note that even for the highest cell seeding densities (and even when viewed on a logarithmic scale), it is difficult to tell by graphical inspection that these data are not exponential. It would be entirely reasonable (and common practice) to fit exponential growth curves to these data to quantitate the growth rates. See the colored lines in Fig. 1. In the following analyses, we fitted exponential growth curves to the synthetic data to simulate widespread practice.

For each synthetic time course dataset (three simulated cell measurements and measurement times for a given replicate), we fitted an exponential growth curve to estimate the replicate’s growth rate. Over all the 60 replicates for each seeding condition, this allowed us to generate a histogram of fitted growth rates (see the inset figures in Fig. 2). For each seeding condition, we used these histograms to generate the mean fitted population growth rate with error bars. See the details in **Methods** and the main part of Fig. 2, where the ground truth of 1/30 hr^-1^ is shown as red lines for comparison. Notice that for any seeding value over 0.1% of the carrying capacity (10^-3^*K*), there was substantial discordance from the ground truth. Indeed, applying a standard statistical test (Student’s *t*-test) for differences in the growth rate between lowest seeding density (10^-5^ *K*) and the other growth rates showed significant (*p* < 0.05) difference between the rates.

**Fig. 2:**
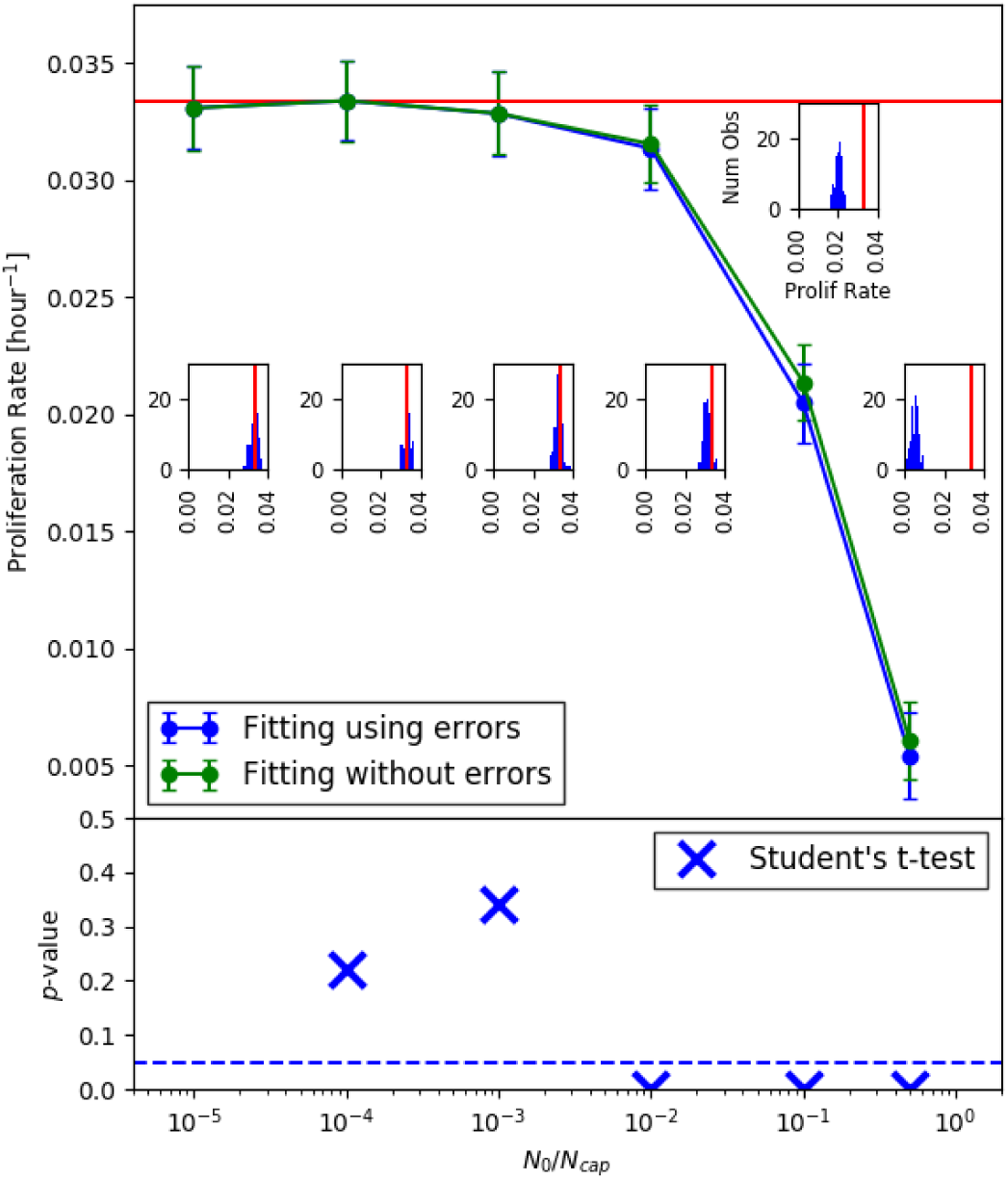
<u>*top and insets:*</u> For initial seeding condition, we fitted exponential growth to each synthetic replicate to get a histogram of the apparent growth rates (insets). As the initial seeding density increased, the deviation between the mean growth rate (green and blue error bars in the main plot) became increasingly discordant from the ground truth rate (red bar). <u>*bottom:*</u> Standard statistical test (Student’s *t-*test) comparing the observed growth rates to the ground truth growth rate showed significant differences (*p* < 0.05) for seeding densities greater than 0.1% of the maximum cell population (10^-3^ *K*).

These differences largely stem from fitting non-exponential growth data to an exponential growth model, which underestimates the growth rate as the cell population approaches its confluent limit. For the standard logistic growth model (see the **Methods** section for the equation), the growth rate decreases from the theoretical maximum value of *α* towards zero as the population increases towards its logistic carrying capacity.

We repeated this synthetic experiment to examine the impact of improving parts of the experimental pipeline, in particular to find experimental improvements could best improve the chances for recognizing when the experimental measurements are experiencing significant confluency effects.

### 2.1 Reducing pipetting error

We first reduced the variability in the initial cell count *N*_0_/*K* to 0.5% (rather than 5%) to simulate improvements in pipetting accuracy, user training, robotic platforms, or other techniques that could potentially be developed. Interestingly, we found that there was little graphical change in the data, as cell counting and measurement timing errors continued to dominate See Fig. 3a.

**Fig. 3.**
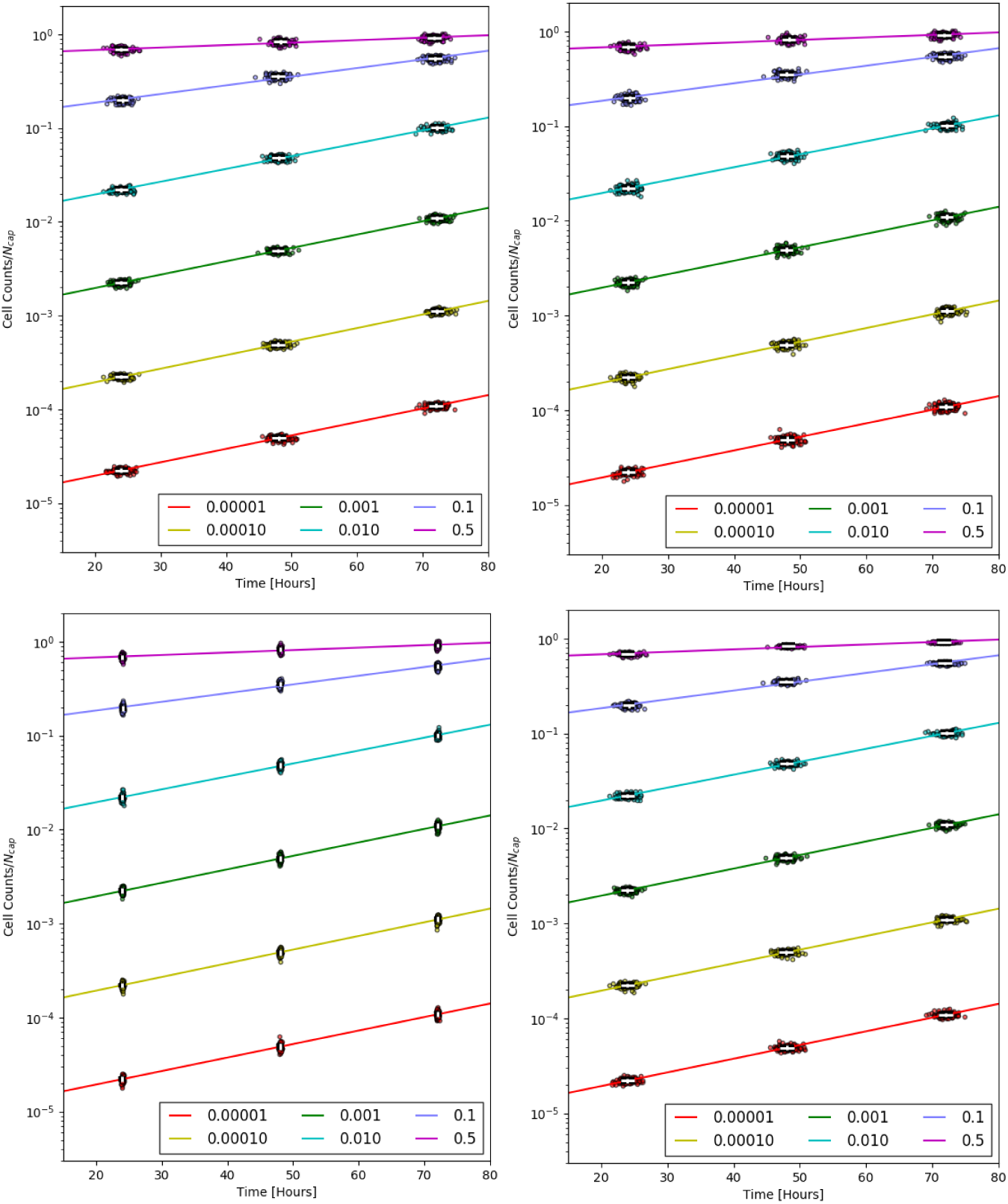
Impact of experimental improvements on the synthetic data. <u>*(top):*</u> Reducing pipetting errors and variability (left) and biological variability (right) had little impact on the overall visual quality, as measurement and timing errors were still dominant. These data would likely still be interpreted as exponential. <u>*(bottom)*</u> Reducing errors in measurement timings (left) and cell counts (right) had a much greater impact on the data appearance; these data have a greater likelihood of being recognized as non-exponential.

### 2.2 Reducing biological variability

Next, we reduced the variability in the growth rates *α* for each biological replicate from 5% to 0.5%, simulating improvements such as robotic-controlled environmental conditions, improved process control (e.g., frequent genetic and proteomic sequencing of feeder cells, molecular profiling of growth media for better consistency), or robotic control of growth medium replacement. As with reducing pipetting error, we found that such theoretical process improvements do little to reduce the apparent noise in the measurements. See Fig. 3b.

### 2.3 Reducing timing errors

We reduced the error in timing the samples to a 0.1 hour standard deviation, to simulate improvements in record-keeping or robotic-controlled imaging. This had a greater effect on the appearance of the data. In particular, we note that the subexponential character of the data for higher seeding densities was much more apparent with reduced timing errors. See Fig. 3c.

### 2.4 Reducing cell count errors

To simulate improvements to image quality and cell segmentation, we simulated the experiment with the measurement errors reduced from 5% to 0.5%. Of all single process improvements, this yielded the greatest improvement to the data. See Fig. 3d.

### 2.5 Reducing all errors

We simulated reducing all the errors (See Sections 2.1-2.4). In this case, the data were much more recognizable as non-exponential. See Fig. 4a.

**Fig. 4.**
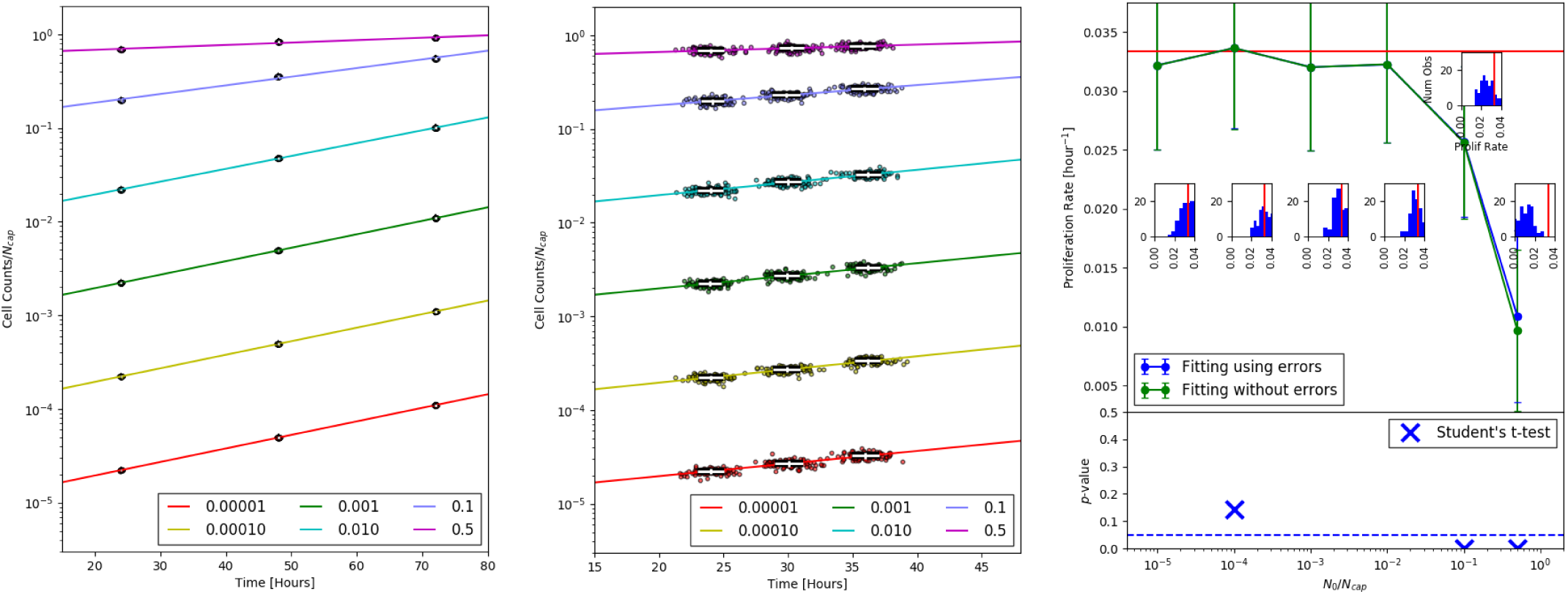
Impact of experimental improvements on the synthetic data. <u>*(left):*</u> If we could reduce all errors and variability by an order of magnitude, it may be more likely to graphically identify the data as non-exponential. <u>(*middle-right*)</u> However, reducing time interval between measurements (middle) reduces the error in the exponential growth approximation when fitting the growth rates, and so even the 1% seeding density comes close to the ground truth (right).

### 2.6 Reducing delays between samples

Lastly, we simulated the same 5% relative errors (and 1 hour timing errors) with more frequent, 12-hour imaging intervals. While this did not improve the overall data appearance, it did improve the concordance of estimated growth rates with the ground truth, because the exponential growth approximation holds better for shorter times. See Figs. 4b-c.

### 2.7 Fitting with errors in time and cell counts.

Examining exponential fits to the synthetic data either taking into account measurement errors in both time and cell counts or just cell count measurement errors, we can see that standard linear regression in logarithmic space produces sufficiently close results to not justify the additional coding expense that we used. See Fig. 2, comparing the two different data points in the top part of the figure.

### 2.8 A computational thought experiment: high-throughput cell phenotyping

These results can also be considered as a computational thought experiment: suppose six different groups were given identical but unlabeled cell lines and asked to profile them in identical conditions (identical oxygenation, growth medium, and culture dishes, but with without specifying the initial seeding density) and compare their results. Would the groups replicate the correct proliferation rates (the ground truth)? And would they correctly identify that they likely had worked with the same cell lines, if they used mismatched cell proliferation rates as a simple rejection criterion?

As we see in Fig. 2, groups with different cell seeding choices could easily observe significantly different (*p* < 0.05) cell proliferation rates. These groups would falsely conclude that they had profiled different cell lines, given the simple rejection criterion. In Figs. 3-4, we see that repeating this exercise with improved instruments, software, and handling consistency would not change the results, due to fitting simple exponential growth models. However, for sufficiently small experimental variability (Fig. 4), the hypothetical groups would be much more likely to determine that they were fitting exponential models to non-exponential data and correct their comparisons. (Note that the *p*-values for the bottom of Fig. 4 exceeded 0.5 and were off the plotted scale for cells seeded at 0.001%, 0.1%, and 1% of the confluent cell population.)

## 3. Discussion

The simulated high-throughput screening experiments demonstrated that the choice of initial cell seeding density can have a substantial impact on the measured cell proliferation rate, even when all the profiled cells are identical. This arises largely from fitting exponential growth models to proliferating cell populations experiencing confluency effects. Mathematical models that do account for confluence effects (e.g., logistic and Gompertzian models) show that proliferation slows from its hypothetical exponential rate even for low cell populations. Biological and technical variability introduced by real experimental systems mask the differences between exponential and sub-exponential growth, making it difficult for experimentalists to graphically recognize the situation and choose more appropriate growth models for fitting.

Our simulations also show that investing resources to reduce pipetting error (Section 2.1), biological variability (Section 2.2) are less likely to “unmask” sub-exponential growth than improvements to measurement timing errors (Section 2.3) or cell counting (Section 2.4). However, even reducing all sources error in the experimental pipeline still yields data that graphically appear exponential (Section 2.5) when sampled three times. However, reducing the delay between measurements (Section 2.6) did reduce the error in estimated proliferation rates, largely because the exponential growth assumption is most valid for small times.

These results have broad implications for implementing, interpreting, and comparing proliferation assays. First, the differences in population growth rates from early growth (far from the confluent limit), mid-term growth (approximately half of the population limit) and growth near the confluent rate are indeed real; proliferation slows as cell populations approach the confluent limit, even without other sources of biological variability. This could help explain the large heterogeneity of measured proliferation rates for commonly used cell lines.

We found that the estimated growth rates were most accurate (closest to the ground truth) for very low seeding densities, on the order of 0.1% of the confluent limit. For high-content screening platforms with growth areas on the order of 0.5 cm^2^ (e.g., the Corning #3094 96-well plate, with 0.32 cm2 growth area [23]), if the confluent cell density is on the order of 10^5^ cells/cm^2^ [24], then such high-throughput systems have confluent cell counts on the order of 10^4^ to 10^5^ cells. If a 0.1% seeding density is necessary for an accurate growth estimate, then we must seed each well with approximately 10 to 100 cells. However, such low seeding populations may not successfully grow. Thus, high-throughput platforms with small well areas will require either (1) cell counting techniques with less than 1% relative error, (2) larger growth areas to accommodate the necessarily low relative seeding density, (3) shorter sampling times to improve the accuracy of the exponential growth assumption when fitting data, or (4) automated fitting to more appropriate growth models that account for confluency effects [18]. Otherwise, such high-throughput profiling experiments may detect confluency differences among different cells, rather than the sought proliferation differences in response to changing drug exposures or other microenvironmental conditions. We do note that shorter sampling times (on the order of 12 hours) allowed our virtual system to estimate the growth rate with less than 5% relative error when seeding at 1% of the confluent limit; but even in this case, seeding at 10% of the confluent limit still gave a relative error of approximately 25%.

For any experimental system (high-throughput or not), it is important to profile cells at a variety of seeding densities, particularly if the confluent limit is unknown. Then the measured growth rates could be plotted against the seeding density (e.g., as in Fig. 2), and the confluent effects would be evident. As a best practice, we would recommend that researchers report the maximum proliferation rate when comparing different cell lines, or the same cell line in different drug or microenvironmental conditions. This way, they can avoid confounding confluency effects. Such practices could help reduce the widespread lack of consensus in finding consistent, “gold standard” ground truths for comparing standard cell lines.

This work highlights the importance of (1) fully recording the experimental design (especially the cell seeding density) and (2) fitting appropriate mathematical models for estimating the population growth rates. Not only will recording the experimental protocols (e.g., using Protocols.io [25]) help to reduce biological and technical variability, but it will also help ensure that we design and apply appropriate analyses to interpret the data. If this allows us to narrow the currently large range in reported cell proliferation and other phenotypic characteristics, then we can improve the quality of curated cell line resources that consistently record the phenotypes of well-characterized, standardized cell lines. This will improve the feasibility and robustness of any high-throughput cell identification platforms, which would seek to match unknown biological samples to the best fit in a library of cell lines. More broadly, we can begin to identify and eliminate the remaining causes of variability in cell line measurements, such as differences in cell growth media, oxygenation, or culture splitting procedures.

Lastly, this works highlights the potential for mathematical modeling beyond typical uses in forming and testing biological hypotheses. Mathematical models of experimental protocols and scientific instruments can help us to better identify the most important sources of error, while assessing where we can most easily make high-impact improvements. In the future, we propose mathematical modeling as an integral part of experimental design.

## 4. Method

The main method creates a simplified digital cell line [6] with two parameters (its birth rate *α* and its confluent density *ρ*_K_, equivalent to a cell carrying capacity *K* on a fixed plate size), and uses analytical solutions to the logistic growth equation as a ground truth for examining the impact of experimental errors and variability. This computational study focuses on estimating the cell population growth rate, and so we leave *K* fixed for the duration of the study. In all this work, *α* is the ground truth. Starting with the ground truth, we create synthetic replicates that simulate the impact of: (1) technical and biological variability in seeding the experimental replicate, (2) variability in the timing of measurements, (3) cell counting errors during measurements, and (4) the limited temporal sampling of an experiment.

### 4.1 Creating a synthetic biological replicate

To account for a biological variability (and to generate a biological replicate), we choose a birth rate *α*\* ~ N(*α*,*ε*), where N(*α*,*ε*) is the normal distribution with mean *α* and standard deviation *ε*. In our work, we set *ε* = 0.05*α* for a 5% relative variability. For greater generality, we could similarly choose the carrying capacity K* ~ N(*K*, 0.05*K*), but we took K* = K for each synthetic replicate to focus our intention on the impact of biological variability in *α*. We set an initial cell population 0 < *N*_*0*_ < K*, which will be varied below to simulate technical variability.

#### Defining the ground truth for a single biological replicate

For any synthetic replicate with actual birth rate *α\** and successfully seeded initial cell population *N*_0_, we model the population growth rate with the standard logistic growth model:

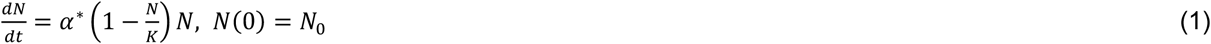

whose analytical solution is

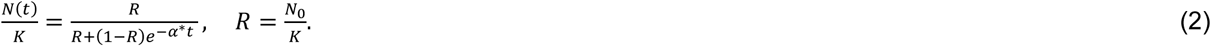

This analytical solution can be evaluated at any time *t* for the *actual* number of cells in the experimental replicate at that time. We then modify this analytical cell count to obtain a measurement (observation) that accounts for other technical errors (see below).

### 4.2 Creating a synthetic technical replicate

For any biological replicate, we create one (or more) technical replicates to simulate technical variability in experiments.

#### Simulating micropipetting and plating

To simulate variability in how many cells are successfully plated during pipetting, we set the initial cell count for each replicate according to 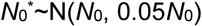, giving a 5% relative variability (standard deviation) in pipetting and plating.

#### Simulating observations with temporal and measurement errors

ATCC [26] and Roche [27] both describe cell proliferation assays as having multiple phases in the following order: lag phase, exponential (logarithmic) growth phase, stationary phase, and death phase. Both methodologies attempt to measure cell growth during the exponential phase by using at least three data points, though many measurements use only two time points (see the growth curves [10] in the NCI Physical Sciences in Oncology Network’s resource of standard operating protocols). To simulate these practices, we choose intended observation times 0 < *T*_1_ < *T*_2_ < *T*_3_; three observations allows us to distinguish between exponential growth (log(*N*) vs. time appears linear) and growth near the confluent limit (log(*N*) vs. time appears sub-linear). Typically for cancer cell culture experiments, *T*_1_ ~ 1-2 days, giving cells time to adhere to the plate and re-enter the cell cycle (i.e., to start measuring past the “lag” phase), and the measurements are on the order of 1 day apart [1, 2, 18]. To simulate timing variability (e.g., an experimentalist needed to wait to access an instrument), we choose 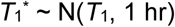, 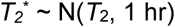, and 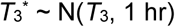.

The simulated “true” cell counts (scaled by *K*) at each of these sampling times are given by:

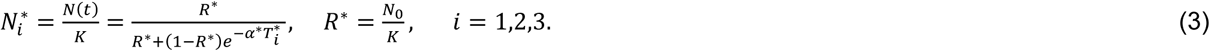

The simulated measurements (observations) 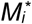 are obtained choosing a single measurement from a normal distribution centered around the true count 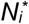 with a relative standard deviation of 5%: 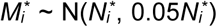 for each observation time 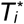. This simulates technical variability in the measurements, such as image segmentation errors.

### 4.3 Putting it all together: a simulated experiment

Each simulated experiment varies the factor *N*_0_/*K* with levels {10^-5^, 10^-4^, 10^-3^, 0.01, 0.1, 0.5}, with a fixed (biological) ground truth *α* and *K*. To simulate a level (with a fixed choice of initial cell count *N*_*0*_, intended sampling times *T*_1_, *T*_2_, *T*_3_, and ground truth *α*, *K*), we simulate 5 biological replicates (each with a different value *α*^*^). For each biological replicate, we simulate 20 technical replicates (each with different values for 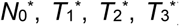), and generate synthetic measurements as indicated above. Thus, each level had 100 total replicates, observed at 3 times, for 300 synthetic measurements. There were six levels of the factor *N*_0_/*K* with values, so each computational experiment generated 1,800 synthetic measurements.

### 4.4 Growth analysis of synthetic data

When fitting the data for each replicate, we used an exponential growth curve on the noisy logistic growth data; this approximates (1) the widespread practice in experimental biology of fitting exponential growth curves, and (2) the general trend that noisy exponential and logistic growth data cannot readily be distinguished graphically. In log space, fitting an exponential curve to data is equivalent to fitting a straight line. Our data though had uncertainties in both time and log(*N/K*), so we used a technique that would allow us to fit in both dimensions [28]. We also performed the same calculation, ignoring the uncertainties in time and log(*N*/*K*), as that only requires the use of traditional linear regression, a more widely known and used technique than that of [28]. We then made histograms of all of the replicates’ growth rates and compared those to the ground truth growth rate (the red vertical line). We also averaged all of the replicates’ growth rates to calculate the “experimental” growth rate over all the replicates.

As part of our analysis, we needed to calculate *p*-values for determining if the “experimental” data could have come from the same distribution as “ground truth” data. We performed a Student’s *t*-test on two distributions of the calculated growth rates done with technique of [28]: One distribution from the lowest seeding density and another distribution from a different seeding density. (Hence, there is no *p*-value for the lowest seeding density).

### 4.5 Numerical details and code availability

All numerical experiments were performed in Python 3.6, using the Anaconda distribution (64-bit Version 4.4.0) [29]. All calculations were done using a fixed random seed to ensure that rerunning the synthetic experiment would result in the same result. All code is available at https://github.com/SamuelFriedman/synthetic_replicates under the BSD 3-clause license.

## Acknowledgments

We thank the Breast Cancer Research Foundation (BCRF) and the Jayne Koskinas Ted Giovanis Foundation for Health and Policy (JKTGF) for the generous support. This work was also funded in part by the National Cancer Institute (1R01CA180149). We thank the Lawrence J. Ellison Institute for Transformative Medicine for prior institutional support.

## References

1. Garvey, C.M., et al., A high-content image-based method for quantitatively studying context-dependent cell population dynamics. Sci Rep, 2016. 6: p. 29752.

2. Patsch, K., et al., Single cell dynamic phenotyping. Sci Rep, 2016. 6: p. 34785.

3. Merouane, A., et al., Automated profiling of individual cell-cell interactions from high-throughput time-lapse imaging microscopy in nanowell grids (TIMING). Bioinformatics, 2015. 31(19): p. 3189–97.

4. Golfier, S., et al., High-throughput cell mechanical phenotyping for label-free titration assays of cytoskeletal modifications. Cytoskeleton (Hoboken), 2017. 74(8): p. 283–296.

5. Gloux, K., et al., Development of high-throughput phenotyping of metagenomic clones from the human gut microbiome for modulation of eukaryotic cell growth. Appl Environ Microbiol, 2007. 73(11): p. 3734–7.

6. Friedman, S.H., et al., MultiCellDS: a standard and a community for sharing multicellular data [preprint]. bioRxiv, 2016: p. 090696.

7. Swat, M.J., et al., Pharmacometrics Markup Language (PharmML): Opening New Perspectives for Model Exchange in Drug Development. CPT Pharmacometrics Syst Pharmacol, 2015. 4(6): p. 316–9.

8. Sluka, J.P., et al., The cell behavior ontology: describing the intrinsic biological behaviors of real and model cells seen as active agents. Bioinformatics, 2014. 30(16): p. 2367–74.

9. DSMZ. MCF-7 Cell line. [cited 2016 June 6]; Available from: https://www.dsmz.de/catalogues/details/culture/ACC-115.html.

10. Physical Sciences in Oncology Network. Background information and SOPs Physical Sciences in Oncology. 2016; Available from: http://physics.cancer.gov/bioresources/SOPs.aspx.

11. National Cancer Institute. Developmental Therapeutic Program NCI-60 Human Cancer Cell Line Screen. [cited 2016 June 6]; Available from: https://dtp.cancer.gov/discovery_development/nci-60/cell_list.htm.

12. Larsson, S., et al., Estimating the Total Rate of DNA Replication Using Branching Processes. Bull. Math. Biol., 2008. 70(8): p. 2177–94.

13. Osborne, C.K., K. Hobbs, and J.M. Trent, Biological differences among MCF-7 human breast cancer cell lines from different laboratories. Breast Cancer Research and Treatment, 1987. 9(2): p. 111–121.

14. Sutherland, R.L., R.E. Hall, and I.W. Taylor, Cell Proliferation Kinetics of MCF-7 Human Mammary Carcinoma Cells in Culture and Effects of Tamoxifen on Exponentially Growing and Plateau-Phase Cells. Cancer Research, 1983. 43(9): p. 3998–4006.

15. Katzenellenbogen, B.S., et al., Proliferation, hormonal responsiveness, and estrogen receptor content of MCF-7 human breast cancer cells grown in the short-term and long-term absence of estrogens. Cancer Res, 1987. 47(16): p. 4355–60.

16. Lepri, S.R., et al., Effects of genistein and daidzein on cell proliferation kinetics in HT29 colon cancer cells: the expression of CTNNBIP1 (beta-catenin), APC (adenomatous polyposis coli) and BIRC5 (survivin). Hum Cell, 2014. 27(2): p. 78–84.

17. Mir Mohammadrezaei, F., et al., Signaling crosstalk of FHIT, CHK2 and p38 in etoposide induced growth inhibition in MCF-7 cells. Cell Signal, 2013. 25(1): p. 126–32.

18. Juarez, E.F., et al., Quantifying differences in cell line population dynamics using CellPD. BMC Syst Biol, 2016. 10(1): p. 92.

19. Macklin, P., When seeing isn't believing: How math can guide our interpretation of measurements and experiments. Cell Systems, 2017 (in press).

20. Poleszczuk, J., P. Hahnfeldt, and H. Enderling, Evolution and phenotypic selection of cancer stem cells. PLoS Comput Biol, 2015. 11(3): p. e1004025.

21. Treloar, K.K., et al., Are in vitro estimates of cell diffusivity and cell proliferation rate sensitive to assay geometry? J Theor Biol, 2014. 356: p. 71–84.

22. Harrison, J.U. and R.E. Baker, The impact of temporal sampling resolutoin on parameter inference for biological transport models. ArXiv e-prints, 2017. 1708: p. 1708.01562.

23. Corning. Corning(r) 96 Well Flat Clear Bottom Black Polystyrene TC-Treated Microplates, 20 per Bag, with Lid, Sterile (Product #3904). 2017; Available from: https://catalog2.corning.com/LifeSciences/enUS/Shopping/ProductDetails.aspx?productid=3904(Lifesciences).

24. Scientific, T.-F. Useful Numbers for Cell Culture. 2017; Available from: https://www.thermofisher.com/us/en/home/references/gibco-cell-culture-basics/cell-culture-protocols/cell-culture-useful-numbers.html.

25. Teytelman, L., et al., Protocols.io: Virtual Communities for Protocol Development and Discussion. PLoS Biol, 2016. 14(8): p. e1002538.

26. American Type Culture Collection. ATCC Animal Cell Culture Guide. 2014; Available from: https://www.atcc.org/~/media/PDFs/Culture%20Guides/AnimCellCulture_Guide.ashx

27. Roche. Culture of Animal Cells - Basic Techniques. 2012; Available from: https://lifescience.roche.com/wcsstore/RASCatalogAssetStore/Articles/Culture%20_of_Animal_Cells-Basic_Techniques_TT.pdf.

28. York, D., et al., Unified equations for the slope, intercept, and standard errors of the best straight line. American Journal of Physics, 2004. 72(3): p. 367–375.

29. Continuum.io. Anaconda (Verison 4.4.0). 2017; Available from: https://www.continuum.io/anaconda-overview.

